# Polygenic risk score of alcohol consumption predicts alcohol-related morbidity and all-cause mortality

**DOI:** 10.1101/652396

**Authors:** Tuomo Kiiskinen, Nina J. Mars, Teemu Palviainen, Jukka Koskela, Pietari Ripatti, Joel T. Rämö, Sanni Ruotsalainen, FinnGen, GSCAN Consortium, Aarno Palotie, Pamela A.F. Madden, Richard J. Rose, Jaakko Kaprio, Veikko Salomaa, Pia Mäkelä, Aki S. Havulinna, Samuli Ripatti

## Abstract

**Objective:** To develop a highly polygenic risk score (PRS) for alcohol consumption and study whether it predicts alcohol-related morbidity and all-cause mortality.

**Design:** Biobank-based prospective cohort study

**Setting:** FinnGen Study (Finland)

**Participants:** 96,499 genotyped participants from the nationwide prospective FinnGen study and 36,499 participants from prospective cohorts (Health 2000, FINRISK, Twin Cohort) with detailed baseline data and up to 25 years of follow-up time.

**Main outcome measures:** Incident alcohol-related morbidity and alcohol-related or all-cause mortality, based on hospitalizations, outpatient specialist care, drug purchases, and death reports.

**Results:** In 96,499 FinnGen participants there were in total 4,785 first-observed incident alcohol-related health events. The PRS of alcohol consumption was associated with alcohol-related morbidity and the risk estimate (hazard ratio, HR) between the highest and lowest quintiles of the PRS was 1.67 [ 95 % confidence interval: 1.52-1.84], p=3.2*10^−27^). In 28,639 participants with comprehensive baseline data from prospective Health 2000 and FINRISK cohorts, 911 incident first alcohol-related events were observed. When adjusted for self-reported alcohol consumption, education, marital status, and gamma-glutamyl transferase blood levels, the risk estimate between the highest and lowest quintiles of the PRS was 1.58 (CI=[1.26-1.99], p=8.2*10^−5^). The PRS was also associated with all-cause mortality with a risk estimate of 1.33 between the highest and lowest quintiles (CI=[1.2-1.47], p=4.5e-08) in the adjusted model. In all 39,695 participants with self-reported alcohol consumption available, a 1 SD increase in the PRS was associated with 11.2 g (=0.93 drinks) higher weekly alcohol consumption (β=11.2 [9.85-12.58 g], p = 2.3*10^−58^).

**Conclusions:** The PRS for alcohol consumption associates for both alcohol-related morbidity and all-cause mortality. These findings underline the importance of heritable factors in alcohol-related behavior and the related health burden. The results highlight how measured genetic risk for an important behavioral risk factor can be used to predict related health outcomes.

## Introduction

Alcohol drinking is a major dose-dependent contributor to morbidity and mortality. Globally, 3 million annual deaths (5 % of all deaths) result from alcohol consumption, and is also linked to more than 200 disease and injury outcomes.(1) As ethanol is a psychoactive substance with addictive properties,(2) alcohol consumption can lead to the development of alcohol use disorders (AUDs), globally prevalent mental disorders of pathological addictive or abusive drinking patterns, which are linked to worse health outcomes, negative socioeconomic effects, and increased mortality.(3) There is a strong connection between the health burden and the level of alcohol consumed, (4) and in total alcohol has been estimated to be the most damaging of all substances of abuse, in terms of harm caused to self and others.(5)

Alcohol-related behaviors are also affected by genetic factors and the estimated heritability of alcohol consumption in twin studies has ranged between 35% and 65% (weighted average 37%) (6) and its single nucleotide polymorphism-based heritability has been estimated to be 10%. (7) Recent large-scale genome-wide association studies (GWAS) have identified multiple loci associated with alcohol consumption, underlining the importance of large study populations for unraveling the genetic architecture underlying alcohol-related traits. (7, 8) Similarly, GWAS of alcohol dependence and the Alcohol Use Disorders Identification Test (AUDIT) scores have shown the traits to be genetically distinct but positively correlated.(9,10)

Polygenic risk scores (PRS) derived from GWAS summary statistics have showcased improved performance in disease prediction. (11) PRSs for known risk factors have also been shown to associate with the related disease (12) and recently associations between multiple risk factor PRSs and related traits were confirmed and reported. (13, 14) However, the link between PRSs for behavioral traits and associated health outcomes remains poorly understood.

The assessment of potential health risks related to alcohol has so far relied on traditional risk factors, including family history, without explicit measurement of genetic risk. Here we developed a highly polygenic risk score for alcohol consumption and studied whether alcohol-related polygenic burden predicts alcohol-use disorders and other alcohol-related morbidity and mortality in Finnish biobank cohorts (n=96,499) linked to electronic health records. Furthermore, we studied whether the PRS for alcohol consumption predicts alcohol-related outcomes beyond self-reported alcohol consumption and other related risk factors, thus providing more objective information independent of individual reporting bias or temporal fluctuations.

## Materials and methods

### Study sample and definition of alcohol-related morbidity

The data is comprised of 96,499 Finnish individuals from FinnGen Data Freeze 2, which includes prospective epidemiological and disease-based cohorts as well as hospital biobank samples (Supplementary Table 1). The data were linked by the unique national personal identification numbers to national hospital discharge, death, and medication reimbursement registries. Additional details and information on the genotyping and imputation are provided in the online-only Supplementary Information.

**Table 1.** Population characteristics of FinnGen, FINRISK, Health 2000, and Twin Cohort-datasets.

|  | FinnGen | FINRISK | Health 2000 | Twin Cohort |
| --- | --- | --- | --- | --- |
| **N** | 96,499<br>(54,262<br>women) | 23,824<br>(12,513) | 5,945<br>(3,260) | 9,926<br>(5,036) |
| **N (incident events)** | 4,785 | 817 | 171 | NA |
| **Age (years)** | 57.5(end of follow-up) | 48.6 (baseline) | 54.0 (baseline) | 49.3 (baseline) |
| **Alcohol drinking (g/week)** | NA | 76.5 | 73.9 | 84.8 |
| **Non-drinkers** | NA | 2,874(12%) | 1,298 (22%) | 30(0.3%) |
| **Current smokers** | NA | 5,929 (25%) | 1,538 (26%) | 3,478 (35%) |
| **Higher education** | NA | 8,612 (36%) | 1,721 (28%) | 1,192 (12%) |
| **Marriage or co-habitation** | NA | 17,468 (73%) | 4,151 (68%) | 6,661 (67%) |
| **GGT (U/I)** | NA | 33.8 | 36.6 | NA |

Alcohol-related baseline measures were available for a subset of the FinnGen dataset consisting of national population survey cohorts: FINRISK, collected in 1992, 1997, 2002, 2007 and 2012 and Health 2000, collected in 2000. The baseline data included self-reported information assessed by questionnaires, anthropometric measures, and blood samples. More detailed descriptions of the FINRISK and Health 2000 studies have been published previously.(15,16)

Additionally, three Finnish twin cohorts, FinnTwin12, NAG-FIN, and Old Twin were pooled and analyzed as one dataset. For these datasets, cohort baseline data was available, but the cohorts were not linked to electronical health records. For details regarding the twin datasets, see the online descriptions (https://wiki.helsinki.fi/display/twineng/Twinstudy). (17,18)

Using nationwide registries for deaths (1969-2016), hospital discharges (1969-2016), outpatient specialist appointments (1998-2016) and drug purchases (1995-2016), we combined 21 somatic and psychiatric alcohol-related diagnoses into a composite disease endpoint, harmonizing the International Classification of Diseases (ICD) revisions 8, 9 and 10, and ATC-codes (**Supplementary Table 1**). These registries spanning decades were electronically linked to the cohort baseline data using the unique national personal identification numbers assigned to all Finnish citizens and residents.

### Genotyping and imputation

FinnGen, FINRISK, Health2000, and Finnish Twin Cohort samples were genotyped with Illumina and Affymetrix genomewide SNP arrays. Individuals with non-European ancestry or uncertain sex were excluded. The details about the genotype calling, quality controls and imputation are provided in the Supplementary Information.

### Polygenic risk scores

Summary statistics from the largest existing GWAS meta-analysis on alcohol consumption (8) were used for constructing the PRS. To avoid overfitting, a separate ad hoc meta-analysis was performed by GSCAN, excluding all Finnish and 23andMe samples (n=527,282 after exclusions). The LDpred-inf method (19) was used to account for linkage disequilibrium (LD) among loci, with whole-genome sequencing data on 2,690 Finns serving as the external LD reference panel. The final scores were generated with PLINK2 (20) by calculating the weighted sum of risk allele dosages for each variant. The number of variants in the final scores was 1.1*10^6^.

### Statistical analysis

The Cox proportional hazard model was used to estimate survival curves, hazard ratios (HRs) and 95 % confidence interval (95% CI) in the survival analyses where age was used as the time scale. R’s cox.zph function was used to test whether the proportional assumption criteria applied in our models. Linear regression in FINRISK and Health 2000 and linear mixed model in the Twin Cohort was used for estimating the relationship between the PRS and alcohol consumption. Logistic regression in the FINRISK and Health 2000 cohorts and linear mixed model in the Twin Cohort was used to estimate the relationship between alcohol abstinence and the PRS.

Age, sex, genotyping array, and the first ten principal components of ancestry were used as core covariates. Additionally, body mass was used as a covariate in the model estimating the PRS-alcohol consumption relationship. Self-reported weekly average alcohol consumption from the past year (when unavailable, the past week’s consumption) was used as the estimate for alcohol consumption. In the fully adjusted survival model analyses, log(x+1)-transformed alcohol consumption-estimate, current smoking status, binary higher education status, binary marital/cohabitation status and GGT (Gamma Glutamyl Transferase) blood levels at baseline served as covariates.

In the survival analyses, all prevalent cases and individuals with covariate missingness were excluded. The PRS was normalized and included as a continuous variable in the models. In the survival analysis the highest and lowest genetic risk for alcohol consumption were compared using PRS quintiles.

In analyses using baseline consumption data, the analyses were performed separately in the Health 2000, FINRISK Study, and Twin Cohorts and then meta-analyzed using fixed effects model.

In risk prediction, FINRISK cohorts with at least 10 years of follow-up (from 1992 to 2002) were used to train the model, and the predictive performance was tested in the Health 2000 cohort. The maximal follow-up window was restricted to 10 years. The change in the predictive performance was assessed by comparing models with and without the PRS using the correlated C-index approach (21) along with calculating the continuous reclassification improvement (NRI) (22) and integrated discrimination improvement (IDI).(23) The Hosmer-Lemeshow goodness-of-fit test was used to test model calibration.

### Patient and public involvement

No patients were involved in setting the research question or the outcome measures, nor were they involved in developing plans for recruitment, design, or implementation of the study. No patients were asked to advise on interpretation or writing up of results. There are no plans to disseminate the results of the research to study participants or the relevant patient community.

## Results

### Cohorts

Our primary dataset (FinnGen) is comprised of 96,499 unrelated individuals (54,262 women) with a total of 55,484,114 person-years of registry-based follow-up and 4,785 first-observed alcohol-related major health events. Alcohol consumption estimates were available for a total of 39,695 individuals from the prospective cohorts (FINRISK, Health 2000 and Twin Cohort). Two cohorts, FINRISK and Health 2000, have full registry data and information on self-reported alcohol consumption and related baseline data and consist of 28,639 individuals (94.5% of the participants after excluding 964 prevalent alcohol-related morbidity cases), with 424,053 person-years of registry-based follow-up and 988 first ever alcohol-related events. The interview-based DSM-IV AUD-status was available in a subset of the Twin cohort for 713 cases and 1460 controls.

### Alcohol consumption

In a meta-analysis of the three cohorts with alcohol consumption estimates available (n=39,888), the PRS for alcohol consumption was strongly associated with self-reported alcohol consumption. A one SD increase in the PRS was associated with an 11.2 g (= 0.93 drinks á 12g) increase in weekly pure alcohol intake (beta=11.2 [9.85-12.6 g], p = 2.3*10^−58^) (**Fig 1, cohort-specific figures in the supplementary material**). Adding the PRS to the model improved r^2^ by ∽0.6 percentage points (from 9.17 % to 9.80 %). In addition, the PRS was negatively associated with alcohol abstinence (reported alcohol consumption 0). In FINRISK and Health2000, a 1 SD increase in the PRS for alcohol consumption was associated with a 13.7% reduced likelihood of being a nondrinker (OR=0.863 [0.833-0.895], p=6.1*10^−16^) while this was not case in the Twin Cohort where there were only 30 nondrinkers (OR=0.999[0.998-1.00] p=0.31). Cohort-specific figures for sex-specific alcohol consumption are in the supplementary material.

**Figure 1.**
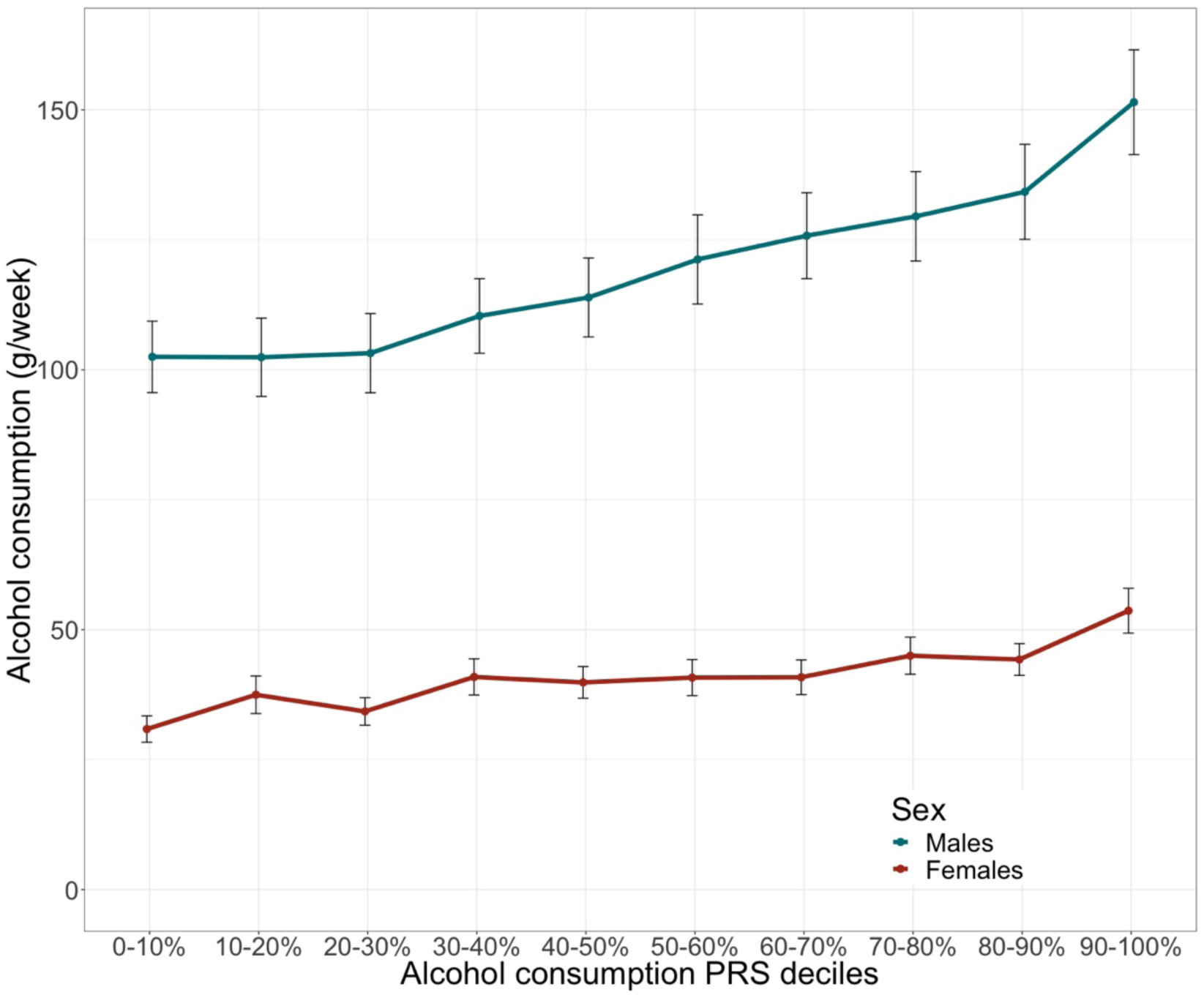
Alcohol drinking (g/week) for the deciles of the alcohol consumption polygenic risk score shown for males (n= 18,887) and females (n= 20,808) with 95% confidence interval error bars (n=39,695)

### Alcohol-related morbidity

The PRS for alcohol consumption was strongly associated with increased risk for lifelong major alcohol-related events derived from electronic health-records in the FinnGen dataset (n=96,499, cases = 4,785) (**Figure 2**). The difference in the risk for alcohol-related morbidity events between the lowest and highest risk quintiles in the PRS was 67 % (HR=1.67 [1.52-1.84], p=3.2*10^−27^) and a 1 SD increase in the PRS was associated with a 21 % increase in risk (HR=1.21 [1.18-1.25], p =1.4*10^−40^). The association was similar in both males (HR=1.21 [1.17-1.26], p=1.8*10^−28^) and females (HR=1.22 [1.16-1-29], p=4.3^10^−14^).

**Figure 2.**
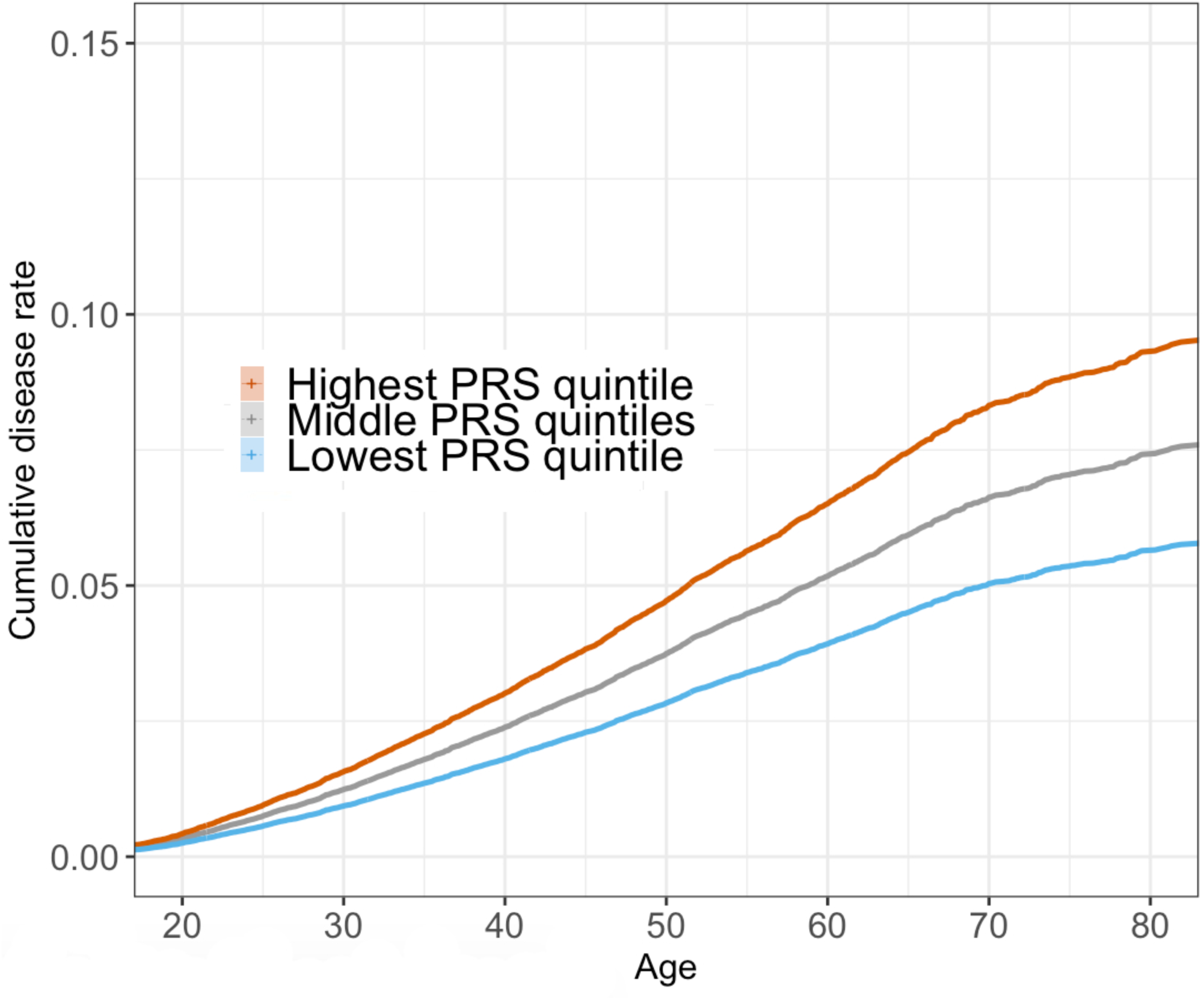
The FinnGen dataset was divided into three groups consisting of the lowest quintile, three middle quintiles and the highest quintile of the alcohol consumption PRS. The cumulative disease rate of alcohol-related morbidity is displayed as a function of age (n=96,499).

In the cohorts where alcohol consumption estimates and other related baseline data were available at the cohort entry time, the PRS was associated with an increased risk of incident major alcohol-related events and the association was maintained also in the fully adjusted model (n = 28,639, cases = 911). In a meta-analysis of the two cohorts 1 PRS SD was associated with a 26% increased risk of incident alcohol-related events when the consumption-estimate was not in the model (HR=1.26 [1.18-1.34], p=1.1*10^−12^) and with a 15 % increase when alcohol consumption was in the model (HR=1.15 [1.08-1.22], p=2.1*10^−5^). In a fully adjusted model, including marital status, education, smoking status and GGT, the estimate were unchanged (HR=1.15 [1.08-1.22], p=2.0*10^−5^) (Table 2).

**Table 2.** Cohort specific and meta-analyzed associations between the alcohol consumption polygenic risk score and alcohol-related **a)** morbidity and **b)** mortality. In the fully adjusted model age, sex, alcohol consumption, smoking, education, marital status and GGT (U/I) were used as non-genetic covariates.

|  | FINRISK | Health 2000 | Meta-analysis |
| --- | --- | --- | --- |
| **a) Alcohol-related morbidity** | **Cases=817** | **Cases=171** | **Cases=988** |
| **Basic model with age and sex** | HR=1.25 [1.16-1.34],<br>p=5.9*10<sup>-10</sup> | HR=1.32 [1.13-1.53],<br>p=0.00036 | HR=1.26 [1.18-1.34],<br>p=1.1*10<sup>-12</sup> |
| **Model with alcohol consumption** | HR=1.13 [1.06-1.21],<br>p=0.00053 | HR=1.23 [1.06-1.43],<br>p=0.0081 | HR=1.15 [1.08-1.22],<br>p=2.1*10<sup>-5</sup> |
| **Fully adjusted model** | HR=1.14 [1.06-1.22],<br>p=0.00027 | HR=1.20 [1.03-1.4],<br>p=0.022 | HR=1.15 [1.08-1.22],<br>p=2.0*10<sup>-5</sup> |
| **b) Alcohol-related mortality** | **Deaths=264** | **Deaths=71** | **Deaths=335** |
| **Basic model with age and sex** | HR=1.21 [1.07-1.37],<br>p=0.0022 | HR=1.41 [1.11-1.8],<br>p=0.0045 | HR=1.25 [1.12-1.4],<br>p=5.9*10<sup>-5</sup> |
| **Model with alcohol consumption** | HR=1.08 [0.952-1.22],<br>p=0.24 | HR=1.34 [1.05-1.71],<br>p=0.017 | HR=1.13 [1.01-1.26],<br>p=0.033 |
| **Fully adjusted model** | HR=1.08 [0.957-1.23],<br>p=0.21 | HR=1.22 [0.965-1.55],<br>p=0.096 | HR=1.11 [0.996-1.24],<br>p=0.058 |

### Mortality

We observed a similar increase in the risk of alcohol-related and all-cause mortality. In FinnGen with 7,249 deaths one SD increase in the PRS for alcohol consumption was associated with 7 % increase in the risk of death (HR= 1.07 [1.05-1.10], p = 4.3*10^−9^). The risk estimate between the highest and lowest 20 % in the PRS was 1.23 (HR = 1.23 [1.14-1.23], p = 1.22*10^−8^). In our prospective cohorts, with cause-of-death information available, 4,125 deaths were recorded (**Table 2**). For all-cause mortality there was 11 % increase in the risk of death per 1 PRS SD in the basic model (HR=1.11[1.07-1.14], p =3.2*10^−10^) and 9 % in the fully adjusted model (HR = 1.09 [1.06-1.12], p=1.1*10^−7^). The risk difference between the highest and lowest quintiles of the PRS was 33 % (HR=1.33 [1.2-1.47], p=4.5e-08) in the fully adjusted model.

Of the 4,125 deaths 335 were known to be alcohol-related. Without alcohol consumption in the model, the increase in alcohol-related mortality was 26 % per 1 PRS SD (HR=1.26 [1.13-1.4], p=3.7*10^−5^). When alcohol consumption was included in the model, the increase was 13 % (HR=1.13 [1.01-1.26], p=0.027) and in a model with all co-variates, 11 % (HR=1.11 [0.996-1.24], p=0.058) (**Table 2**). Similarly, the PRS was associated with a higher risk of death from other than alcohol-related causes (n=3,790) when fully adjusted for all covariates (HR=1.08 [1.05-1.12], p=1.4*10^−6^).

### DSM-IV Alcohol-use disorder

The PRS was also associated with an interview-based DSM-IV alcohol use disorder diagnosis in the Nicotine Addiction Genetics Family cohort (440 cases, 1,140 controls) and a subset of FinnTwin16 cohort (273 cases, 320 controls). A meta-analysis of the two cohorts (713 cases) resulted in a combined 20 % increase in the prevalence of AUD per 1 PRS SD (OR = 1.20 [1.11-1.31], p = 2.29*10^−5^) in the unadjusted model. Adjusting for marital status, education and smoking explained part of the effect (OR=1.14 [1.02-1.28], p=0.023) and further adjusting with maximal amount of drinks taken explained most of the effect (OR=1.06 [0.94-1.19], p=0.35).

### Prediction

The predictive performance of the PRS was evaluated in the Health 2000 cohort (5,732 complete cases, 110 events) with a follow-up-window of 10 years based on the Cox model trained in the FINRISK cohort (18427 complete cases with ≥10 years of follow-up, 628 events). In a model not including the alcohol consumption estimate, adding the PRS to the model increased the C-index by 0.020, from 0.69 to 0.71 (p = 0.017). Both IDI (0.00242 [0.00102-0.00383], p = 7.3*10^−4^) and NRI (0.335 [0.146-0.523], p = 5.1*10^−3^) shifts were positive and statistically significant. When the log-transformed alcohol consumption estimate was included, a modest improvement of prediction was observed (C-index=0.0022 from 0.812 to 0.814, p-value=0.30; NRI=0.308 [0.119-0.497], p = 0.0014 and IDI=0.00173 [0.000726-0.00305], p = 0.017). Similarly, a modest gain was observed when adding PRS to a model with all available covariates including also marital status, education status, smoking status and GGT (C-index=0.00183 from 0.847 to 0.849, p=0.44; NRI=0.235 [0.0461-0.423], p=0.015; IDI =0.00331[0.0000254-0.00659], p = 0.048).

## Discussion

We developed a highly polygenic risk score for alcohol consumption by obtaining weights from a recently published large-scale discovery sample and showed that the PRS was strongly associated with alcohol consumption in independent biobank cohort samples. An increased polygenic burden for alcohol consumption was associated with higher incidence of major alcohol-induced health events. The associations remained significant when we accounted for self-reported alcohol consumption and other relevant covariates; in a fully adjusted model the relative risk-estimate between the highest and lowest quintiles of the polygenic risk score was 1.6. Furthermore, the PRS was also associated with both alcohol-related, non-alcohol related and all-cause mortality.

### Comparison with other studies

Our PRS shows the utility of genetic information for prediction of alcohol-related harm. The PRS, developed from a genetic analysis of cross-sectional self-reported alcohol consumption, was associated with future risk of major alcohol-related health events. While a large number of PRSs have already been established for various traits and diseases (11), the development of PRSs for behavioral traits, such as substance use, has until now been limited (24-27) and the studies have not assessed their impact on future major health events.

### Implications

Our results show that using a large sample size with long follow-up, we were able to build a PRS of alcohol consumption that is associated not only with alcohol consumption in independent samples, but also with future incident alcohol-related health events. In line with the knowledge that alcohol consumption is a major contributor to the worldwide burden of death, especially among working-age adults (1), we found the PRS to be associated also with all-cause mortality, further highlighting the importance of alcohol drinking as a cause of premature death.

Our score provides a genetic basis for potentially identifying a subset of high-risk individuals even early on in life, with potential for more targeted prevention of AUDs and other alcohol-related morbidity. Prevention is a cost-effective and efficient strategy to reduce alcohol related harms (28) and it is labeled one of the United Nations main health-related worldwide strategies of sustainable development (https://sustainabledevelopment.un.org/sdg3). A higher genetic predisposition for alcohol-related harms was detected both in the presence and absence of alcohol consumption data, as our PRS predicted alcohol-related harms beyond self-reported alcohol consumption. Health services are encouraged to support initiatives for screening and brief interventions for harmful drinking (29) as an effective strategy for tackling alcohol-related harm.(30) Thus, genetic information could potentially be used to improve the arsenal of possible strategies to detect high-risk individuals for targets of brief interventions. The fact that individuals in the highest PRS quintile showed an elevated risk for alcohol-related health events even in fully adjusted models could justify the use of genetic information even in clinical settings where a detailed history of alcohol consumption estimates, AUDIT-scores, or similar information are attainable. Communicating the information of higher risk for alcohol-related harm to patients could serve as a motivator for reducing drinking or committing to abstinence. The effect of being aware of one’s negative expectations could also be unwanted, as is thought to be the case in the stereotype threat phenomenon. (31-34)

Self-reported alcohol consumption is known to be biased and problematic in terms of reliability and validity for predicting alcohol-related risks (35,36). Some bias derives from true measurement error, but another source is the lifelong temporal fluctuation of alcohol-drinking patterns not captured by a measure at one single timepoint. Our PRS was associated with alcohol-related harms even when adjusting for self-reported alcohol consumption estimate. One potential reason for this is that the PRS contains information from the latent genetic predisposition for alcohol consumption, thus overriding both the true measurement error and temporal fluctuations in alcohol drinking volume.

It has been hypothesized that alcohol consumption-based genetic discovery might inform more about low-level drinking than about problematic drinking and AUDs.(37) However, we built a polygenic risk score for alcohol consumption and successfully used it to predict alcohol-related harms. Due to the robustness of a self-reported single timepoint alcohol consumption estimate and the fact that different alcohol-related traits are to some degree genetically distinct,(9,10) it is expected that a PRS developed directly for alcohol-related morbidity will outperform our PRS in predicting alcohol-related health burden. Supporting this assumption, the general pattern is that PRSs are more strongly associated with their respective diseases than with related phenotypes. (13,14) Unfortunately, no high-quality summary statistics for alcohol-related harms including both somatic and psychiatric outcomes yet exist; the performed GWAS have only covered alcohol dependence (10) and been far smaller in size than our discovery sample of choice, thus making future efforts for large-scale GWAS discovery based on alcohol-related harms more than necessary.

### Strengths and limitations

Our polygenic risk score was derived using European ancestry discovery samples and tested in the Finnish population. Its applicability in other populations therefore needs further evaluation as the alcohol-related genetic mechanisms may vary between populations. However, it has to be noted that the PRS derived from a non-Finnish sample performed well in the Finnish dataset, even though Finns are somewhat genetically different from the rest of the Europeans (38).

Our design allowed us to study outcomes prospectively. Our registry-based follow-up captures alcohol-related outpatient and inpatient visits, withdrawal treatment prescription for alcoholism, and deaths, thus covering major alcohol-related health events over several decades. Nonetheless, some of the milder cases of alcohol-related health problems could have gone undetected.

### Conclusions

In conclusion, a polygenic risk score for alcohol consumption was associated with elevated risk for incident alcohol-related health events and all-cause mortality. These findings underline the importance of heritable factors driving alcohol-related behavior. A successful attempt to predict alcohol-related health outcomes with a polygenic risk score shows promise in possible future utilization of genetic information in risk estimation and prediction of alcohol-related harms.

## Footnotes

### Contributors

All authors contributed to the study concept and design, analysis and interpretation of the data, as well as to the critical revision of the manuscript for important intellectual content or additionally to data acquisition. GSCAN provided the GWAS summary statistics. TK, TP, NJM and JKa performed the statistical analyses with support from SRi, ASH, JKo and SRu. ASH connected the data to the registries. TK drafted the manuscript. The corresponding author attests that all listed authors meet authorship criteria and that no others meeting the criteria have been omitted.

### Ethical approval

The study was approved by the Coordinating Ethical Committee of the Helsinki and Uusimaa Hospital District, reference numbers HUS/990/2017 (FinnGen), 246/13/03/00/15, 113/E3/2001 and HUS/1169/2016 (The Twin Cohort). The FINRISK and Health 2000 data are stored in the THL Biobank. The transfer of the FINRISK and Health 2000 sample collections to the THL biobank has been approved by the Coordinating Ethics Committee of Helsinki University Hospital on 10^th^ October 2014 and by the Ministry of Social Affairs and Health on 9^th^ March 2015. This study was conducted under the THL biobank permission BB2017_64 (FINRISK and Health 2000). No additional ethical approval was needed for meta-analysing the results.

## Acknowledgements

We would like to Lea Urpa for proofreading, and Sari Kivikko, Huei-Yi Shen and Ulla Tuomainen for management assistance. We thank all participants of the study cohorts for their generous participation.

