## Supplementary material for "Polygenic risk score of alcohol consumption predicts alcohol-related morbidity and all-cause mortality"

**Supplementary methods**

**Genotyping and imputation**

FinnGen samples were genotyped with Illumina and Affymetrix arrays (Thermo Fisher

Scientific, Santa Clara, CA, USA). Genotype calls were made with GenCall and zCall algorithms for

Illumina and AxiomGT1 algorithm for Affymetrix chip genotyping data. Genotyping data produced

with previous chip platforms were lifted over to build version 38 (GRCh38/hg38) following the

protocol described here: dx.doi.org/10.17504/protocols.io.nqtddwn. Samples with sex discrepancies, high genotype missingness (>5%), excess heterozygosity (+-4SD) and non-Finnish ancestry were removed. Variants with high missingness (>2%), deviation from HWE (P <1e-6) and low minor allele count (MAC<3) were removed. Pre-phasing of genotyped data was performed with Eagle 2.3.5 (https://data.broadinstitute.org/alkesgroup/Eagle/) with the default parameters, except the number of conditioning haplotypes was set to 20,000. Imputation was carried out by using the populationspecific SISu v3 imputation reference panel with Beagle 4.1 (version 08Jun17.d8b, https://faculty.washington.edu/browning/beagle/b4_1.html) as described in the following protocol: dx.doi.org/10.17504/protocols.io.nmndc5e. SISu v3 imputation reference panel was developed using the high-coverage (25-30x) whole genome sequencing data generated at the Broad Institute of MIT and Harvard and at the McDonnell Genome Institute at Washington University; and jointly processed at the Broad Institute. Variant callset was produced with GATK HaplotypeCaller algorithm by following GATK best-practices for variant calling. Genotype-, sample- and variant-wise QC was applied in an iterative manner by using the Hail framework v0.1 (https://github.com/hail-is/hail). The resulting high-quality WGS data for 3,775 individuals were phased with Eagle 2.3.5 as described above. Post-imputation quality control involved excluding variants with INFO score < 0.7.

The FINRISK and Health 2000 samples were genotyped using Illumina CoreExome, OMNIExpress, and 610K arrays. Individuals with non-European ancestry or obscure sex were excluded. Quality control (QC) before phasing and imputation excluded variants with missingness >5%, call rate <95%, minor allele count (MAC) <3 (if Zcalled) or MAC <10 (if called using Illumina GenCal), INFO <0.8, minor allele frequency <0.001%, Hardy-Weinberg equilibrium p-value <1*10-10, and heterozygosity exceeding ±4 standard deviations. The QC was performed on simultaneously on all data. Prior to imputation, the haplotypes were estimated using SHAPEIT2. (1) Imputation was done with IMPUTE2 (2) using high-coverage, population-specific reference panels of 2690 whole-genome and 5093 whole-exome sequences.

Genotyping of the twin cohorts was done using Illumina Human610-Quad v1.0 B, Human670-QuadCustom v1.0 A, Illumina HumanCoreExome- (12 v1.0 A, 12 v1.1 A, 24 v1.0 A, 24 v1.1 A, 24 v1.2 A) and Affymetrix FinnGen Axiom arrays. The algorithm for genotype calling were Illumina’s GenCall for all HumanCoreExome chip genotypes, Illuminus for 610k & 670k chip genotypes and AxiomGT1 for Affymetrix chip genotypes. Genotype quality control were done in three batches (batch1: 610k+670k, batch2: HumanCoreExome and batch3: Affymetrix chip genotypes) with removing variants with call rate below 97,5% (batch1 and batch3) and 95% (batch2), removing samples with call rate below 98% (batch1) or 95% (batch2 and batch3), removing variants with its minor allele frequency below 1% and Hardy-Weinberg Equilibrium p-value lower than 1e-06. Also, samples from all batches with heterozygosity test method-of-moments F coefficient estimate value below -0.03 or higher than 0.05 (batch1 and batch2) or ±4SD from the mean (batch3) were removed along with the samples which failed sex check or were among the MDS principal component analysis outliers. Total amount of genotyped autosomal variants after QC were 475526 (batch1), 239894 (batch2) and 388673 (batch3). We then performed pre-phasing using Eagle v2.3 (3) and imputation with Minimac3 v2.0.1 using University of Michigan Imputation Server. (4) Genotypes of all batches were imputed to Haplotype Reference Consortium release 1.1 reference panel. (5)

**Supplementary figures and tables**

**Supplementary Figure 1.** *Sex-specific self-reported alcohol-consumption estimate distribution in* ***a)*** *FINRISK (the smaller inserted graph in the right upper corner of the picture zooms into lower volumes of alcohol consumption thus providing better resolution)* ***b)*** *Health 2000 and* ***c)*** *combined twin population survey cohorts.*

**a)**

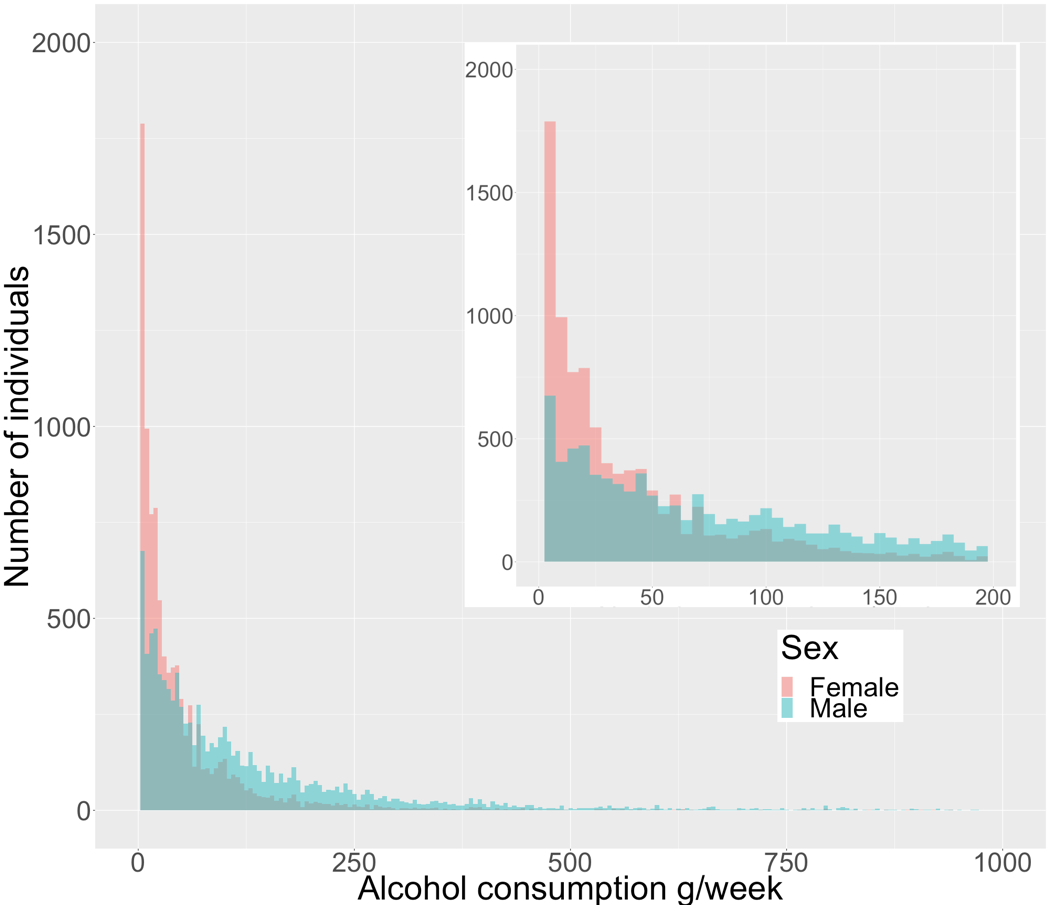

**b)**

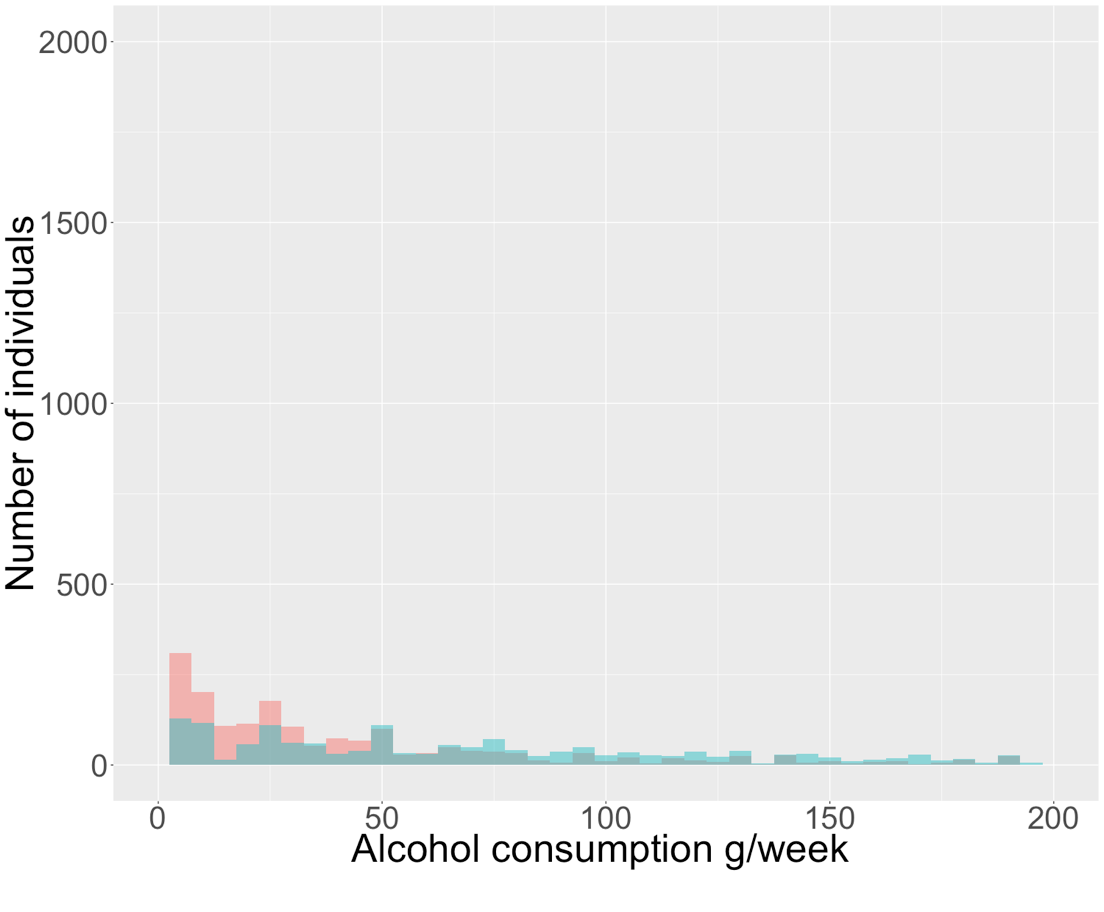

**c)**

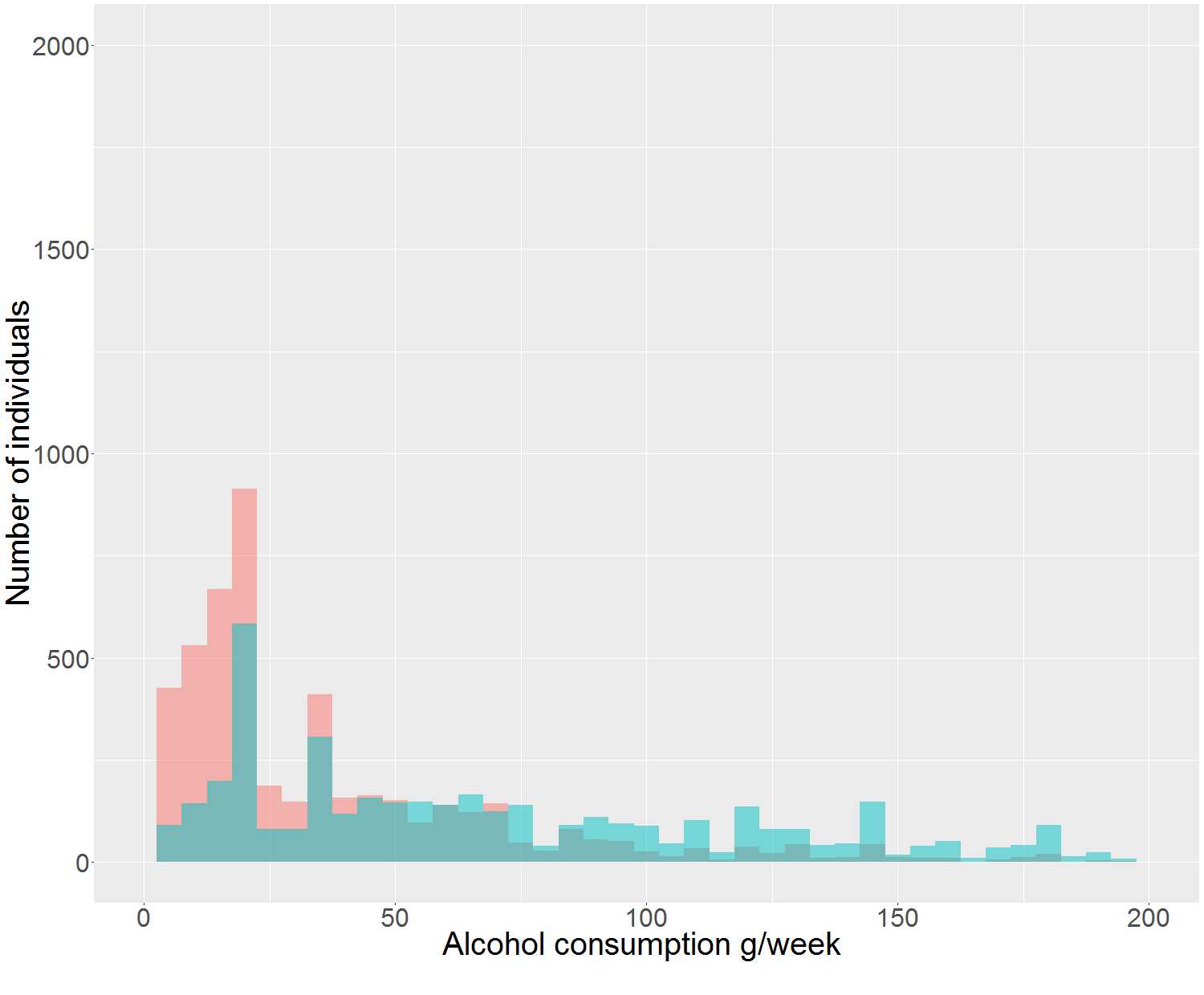

**Supplementary Figure 2.** *Cohort specific alcohol drinking (g/week) for the deciles of the alcohol consumption PRS shown for males and females with 95% confidence interval error bars in* ***a)*** *FINRISK* ***b)*** *Health 2000 and* ***c)*** *Finnish Twin cohorts.*

**a)**

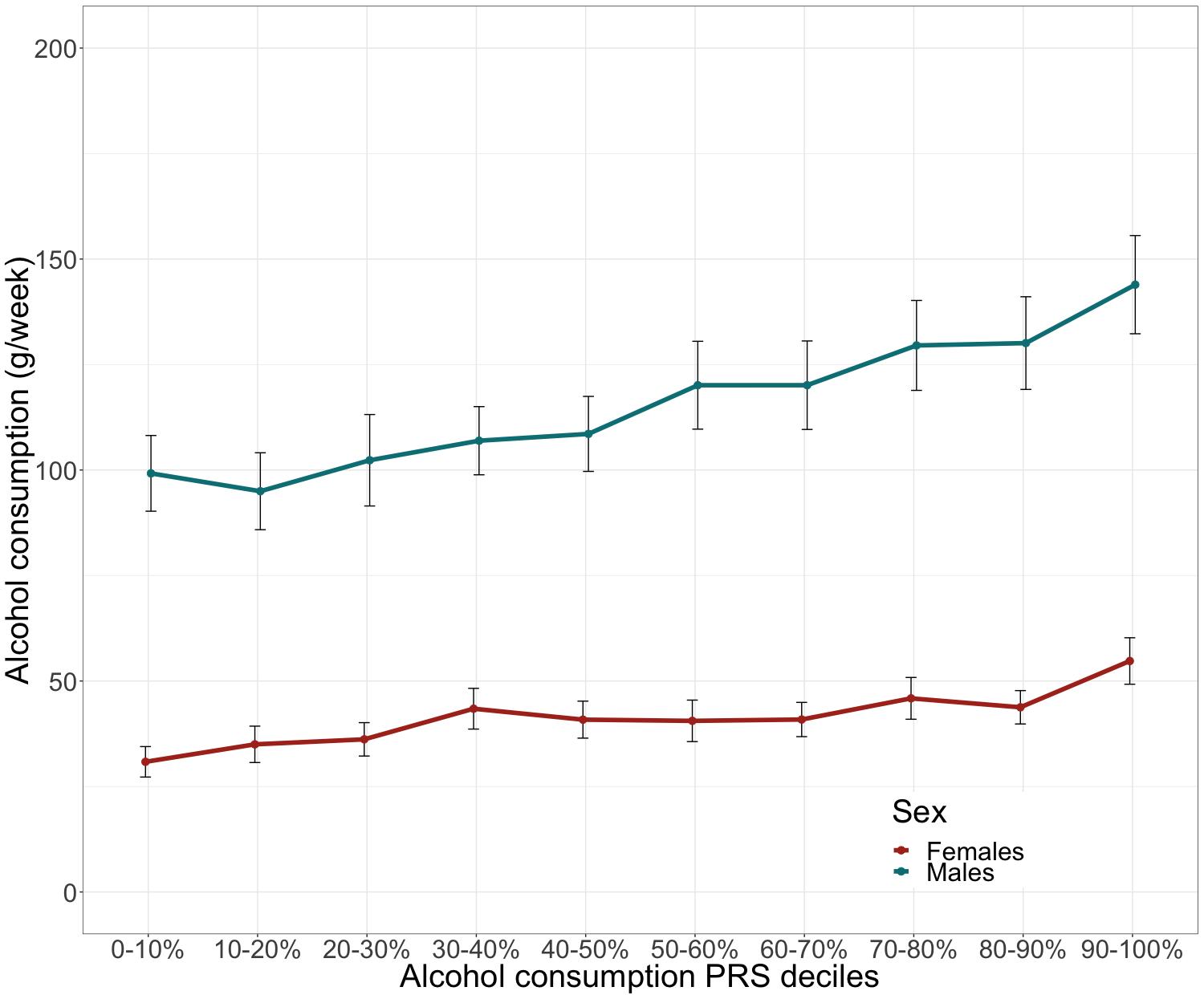

**b)**

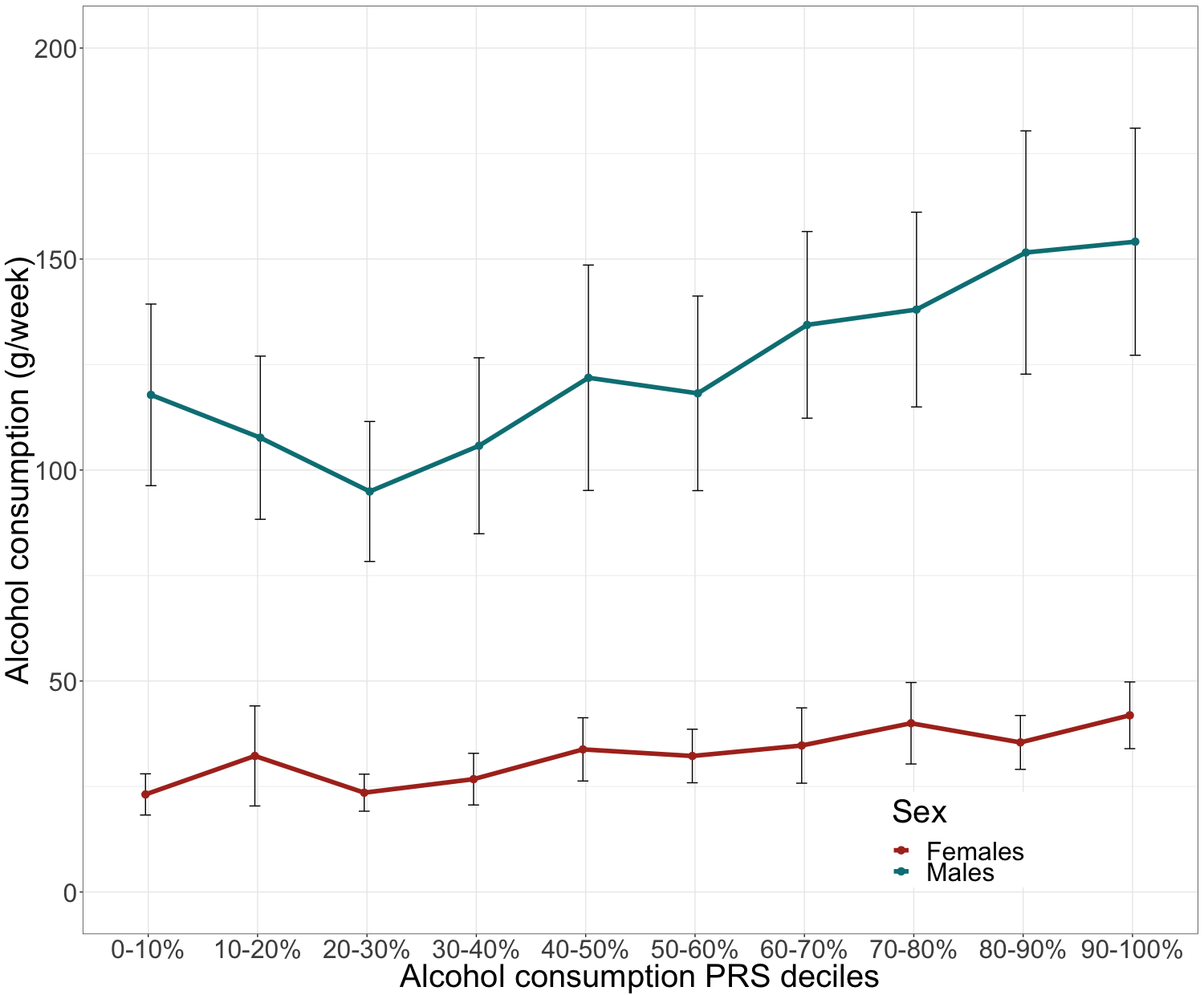

**c)**

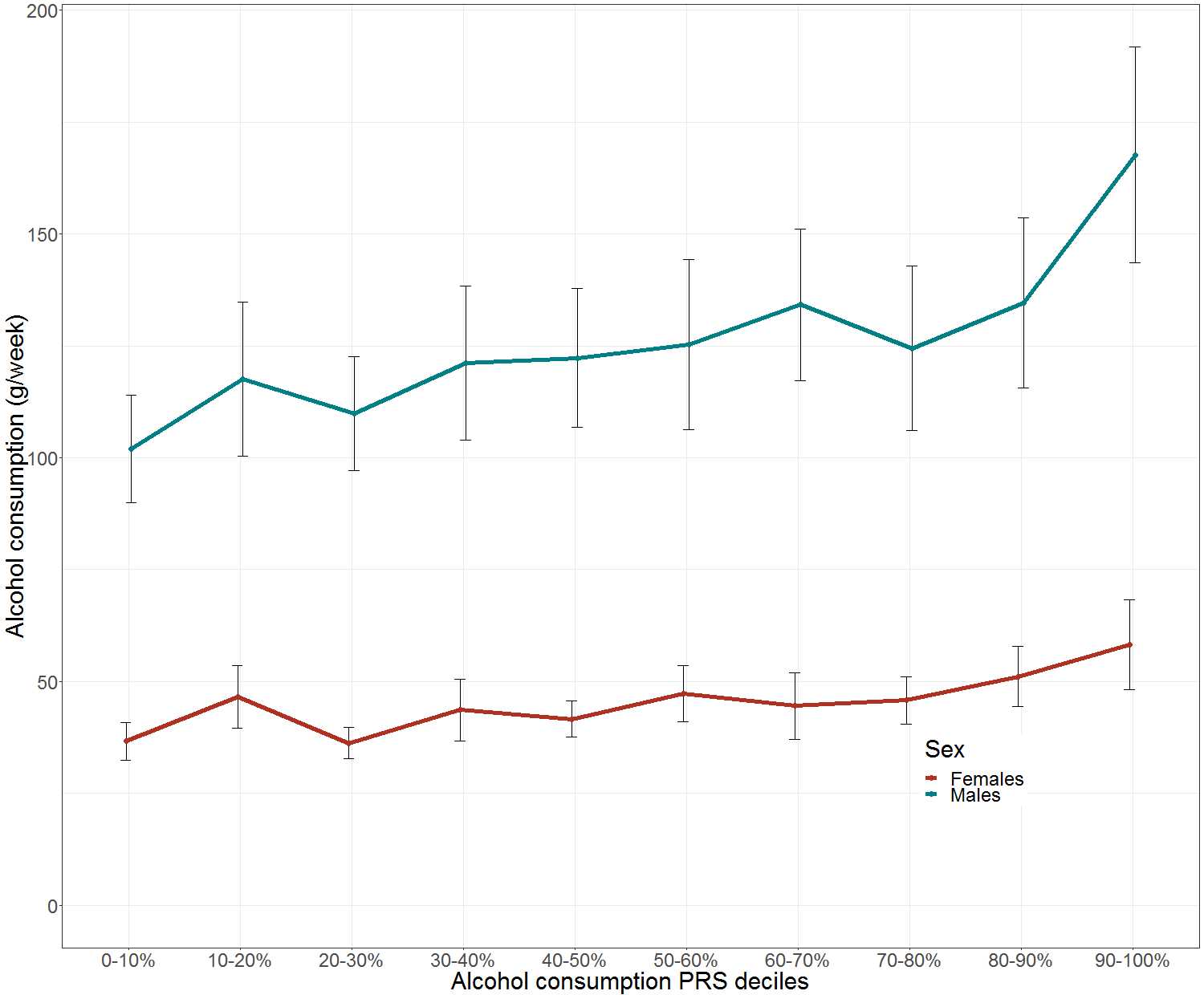

**Supplementary Table 1.** *Specific conditions and corresponding ICD/ATC codes that were used in the construction of the combinatory alcohol-related morbidities endpoint. As ICD-10 F10.0 diagnosis can be used only in the absence of AUDs, we included it only in the alcohol-related mortality endpoint definition.*

|  | ICD-10 | ICD-9 | ICD-8 | ATC |
| --- | --- | --- | --- | --- |
| Acute alcohol intoxication* | F10.0 |  |  |  |
| Mental and behavioural disorders due to alcohol, excluding non-pathological acute intoxication | F10.1-9 | 291,303,305A | 291,303 |  |
| Degeneration of nervous system due to alcohol | G31.2 |  |  |  |
| Epileptic seizures related to alcohol | G41.51 |  |  |  |
| Alcohol induced polyneuropathy | G62.1 | 3575A |  |  |
| Alcoholic myopathy | G72.1 |  |  |  |
| Alcoholic cardiomyopathy | I42.6 | 4255 |  |  |
| Maternal care for (suspected) damage to fetus from alcohol | O35.4 |  |  |  |
| Alcoholic gastritis | K29.3 | 5353A |  |  |
| Alcoholic liver disease | K70 | 5710-3 | 5710 |  |
| Acohol-induced acute pancreatitis | K85.2 | 5770D-F |  |  |
| Alcohol-induced chronic pancreatitis | K86.0 | 5771C-D |  |  |
| Fetus and newborn affected by maternal use of alcohol | P04.3 | 7607A |  |  |
| Accidental poisoning by and exposure to alcohol | X45 |  |  |  |
| Guidance and medical advice to a person with alcohol abuse | Z71.4 |  |  |  |
| Alcohol-induced pseudo-Cushing syndrome | E24.4 |  |  |  |
| Toxic effect of ethanol |  |  |  |  |
| Toxic effect of unspecified or or other (than ethanol) alcohols | T51.1-9 | 9801-9 | 9801-9 |  |
| Use of disulfiram, acamprosate or naltrexone |  |  |  | N07BB01, N07BB02, N07BB04 |

**Supplementary Table 2.** *The prospective epidemiological and disease-based cohorts and hospital-based samples in FinnGen Data Freeze 2*

| Cohort | N |
| --- | --- |
| Auria biobank* | 3,556 |
| Blood Service biobank | 6,271 |
| Borealis biobank* | 1,383 |
| Botnia Family1 | 1,176 |
| Botnia New | 6 |
| Botnia PPP4 | 4,724 |
| Botnia Sib-Helsinki | 428 |
| FinHealth 2017 | 5,938 |
| FINRISK 1992-2012 | 29,774 |
| GeneRISK | 7,090 |
| Health 2000 | 6,672 |
| Helsinki biobank* | 10,410 |
| Health 2011 | 717 |
| Migraine | 7,882 |
| SUPER | 4,420 |
| Diabetes | 6,052 |
| Sum | **96,499** |

*Hospital-based samples
